# Elucidating Resistance Mechanisms in *Staphylococcus epidermidis*: A High-Performing MALDI-TOF MS-Based Proteomic Approach for Predictive Modeling

**DOI:** 10.1101/2024.01.16.575876

**Authors:** Michael Ren, Qiang Chen, Jing Zhang

## Abstract

The emergence of *Staphylococcus epidermidis* as a significant nosocomial pathogen necessitates advancements in more efficient antimicrobial resistance (AMR) profiling. However, existing culture-based antimicrobial susceptibility testing (AST) methods can take up to 96 hours, while holistic PCR assays can cost thousands of dollars. This study combines machine learning with matrix-assisted laser desorption/ionization time-of-flight mass spectrometry (MALDI-TOF MS) to develop predictive models for various antibiotics using a comprehensive dataset containing thousands of *S. epidermidis* isolates. Optimized machine learning models utilized feature selection and achieved high AUROC scores ranging from 0.80 to 0.95 while maintaining AUPRC scores up to 0.96 for balanced datasets. Shapley Additive exPlanations (SHAP) were employed to analyze relevant features and assess the significance of corresponding protein biomarkers while also verifying that predictive power was derived from the detection of proteins rather than noise. AMR prediction models were validated externally to evaluate model performance outside the original data collection site. The findings in this study demonstrate that combining machine learning with AMR profiling yields significant and relevant results, indicating that such an approach is a promising solution for rapid and cost-effective treatments for nosocomial infections. Models created to predict AMR can also aid in biomarker discovery. The workflow used in the present study is potentially applicable to other microbial pathogens in the future.

## 1. Introduction

Matrix-assisted laser desorption/ionization time-of-flight mass spectrometry (MALDI-TOF MS) is an emerging technique that has solidified its usefulness in routine and rapid microbial species identification (Candela et al., 2022). Despite its practicality, the implementation of MALDI-TOF MS in antimicrobial susceptibility testing (AST) has yet to be fully explored. Previous research attempted to predict resistance based on the existence of singular peaks, but such methods are inconsistent and do not consider significant patterns of multiple peaks that impart resistance in combination (Wolters et al., 2011; Wang et al., 2022). More recent works have demonstrated the potential of artificial intelligence (AI) in detecting such patterns and successfully predicting resistances for clinically relevant bacteria such as *Staphylococcus aureus*, *Escherichia coli*, *Klebsiella pneumoniae*, and *Enterococcus faecium* (Weis et al., 2022; Wang et al., 2022). However, *Staphylococcus epidermidis*, a member of the coagulase-negative staphylococci (CoNS) bacterial family, has recently emerged as a formidable pathogen, gaining prominence as a leading agent behind nosocomial infections within the body (CDC, 2004). To date, no studies have demonstrated a substantial MALDI-TOF MS approach for antimicrobial resistance (AMR) prediction in *S. epidermidis*.

Under normal circumstances, *S. epidermidis* is a standard component of human epithelial microflora and maintains a harmless relationship with humans. However, many *S. epidermidis* isolates can form biofilms—conglomerates of multiple cells that form on surfaces—and pose an issue due to the difficulty of eradicating such formations. Due to the ubiquity of *S. epidermidis* on various surfaces, *S. epidermidis* has become a common contaminant of medical devices despite the sanitation protocols typically implemented within hospital settings, leading to various implications (Costerton et al., 1999). For example, *S. epidermidis* stands out as the predominant cause of infections associated with indwelling medical devices. One study from the United States indicated that at least 22% of central intravenous catheter insertions were caused by *S. epidermidis* alone (O’Grady et al., 2002). A different study found that *S. epidermidis* was behind approximately 13% of prosthetic valve endocarditis infections, which was followed by an intracardiac abscess rate and mortality rate of 38% and 24%, respectively (Chu et al., 2009). Such infections are of serious concern because *S. epidermidis* is extremely difficult to treat once developed, which incurs annual expenditures of more than $2 billion in the United States healthcare system alone (Otto, 2009). The difficulty in treating these infections lies in their intrinsic resistance to antibiotics and host defense systems due to biofilm formation (Costerton et al., 1999).

As such, it is imperative to develop more rapid and effective methods of diagnosis for *S. epidermidis* to treat infections before they can progress. By facilitating the early detection of microbes and gaining an in-depth understanding of their inherent resistance mechanisms, healthcare professionals can significantly improve the precision and effectiveness of their treatments (Huang et al., 2013). This necessitates extensive AMR profiles to ensure optimal treatment plans are prescribed and tailored to the specific microbe at hand. These plans benefit not only the patient but also contribute to antibiotic stewardship efforts to combat the rise of multidrug-resistant bacteria (CDC, 2019).

However, existing methods for AMR profile creation come with many drawbacks that make diagnosis suboptimal, thereby increasing patient morbidity, mortality, and healthcare costs (Otto, 2009). Resistance to antibiotics is commonly determined by culture-based methods, which are known to have significant drawbacks. For one, it can take up to 96 hours for a complete AMR resistance profile to be reported, representing a significant time gap between sample collection and results (Banerjee et al., 2021). In addition, these methods are expensive and require a high level of technical expertise to ensure that AMR assays are completed successfully (McLain et al., 2016). Using culture-based methods can, therefore, delay the delivery of treatment decisions, which leads to the prescription of suboptimal antibiotics—either too narrow- or broad-spectrum—that can complicate treatment and put the patient’s health at risk (Banerjee et al., 2015; Kommedal et al., 2016).

Another approach to determining antibiotic resistance entails the process of amplifying genes and their subsequent analysis via polymerase chain reactions (PCR). Although this method provides results in up to 48 hours, which is significantly faster than culture-based methods, the economic burden is extremely high. A holistic resistance assay of multiple antibiotics with PCR can cost approximately $4,900 (Aggarwal et al., 2020). PCR is generally performed on single, targeted genes, which prevents the diagnosis from considering other factors, including gene specificity and resistance that is not genetically mediated. The complexity of these procedures increases significantly when comprehensive AMR profiles are necessitated, as susceptibility testing for each antibiotic requires an AMR assay independent from the others, which only accentuates the inherent limitations associated with their implementation and accounts for the high costs of holistic profiling (Weis et al., 2022).

As such, looking towards other unexplored methods like MALDI-TOF MS to predict AMR is essential. MALDI-TOF-MS works by profiling bacterial extracts and constructing a mass spectral fingerprint of microorganisms, which can be subsequently matched with databases of known mass spectra patterns (Carbonnelle et al., 2011). Recent studies have suggested that MALDI-TOF MS could also serve as a potential tool for AMR profiling (Weis et al., 2022; Wang et al., 2022). This can be accomplished by extracting additional information and feature combinations from MALDI-TOF MS data that is likely to contain biomarkers associated with resistance to a variety of antibiotics. Moreover, MALDI-TOF MS has proven to be advantageous over traditional AMR assays due to its speed, precision, and low cost. Studies have found that the diagnostic turnaround time for MALDI-TOF MS is approximately 38 minutes and that MALDI-TOF MS takes a mere $0.50 per analysis (Dhiman et al., 2011). As a result of these advantages, MALDI-TOF MS has become the most widespread method for species-level microbial identification in clinical laboratories, which means that efficient integration of the machine learning workflow for AMR prediction into existing sites would be feasible (Weis et al., 2022). Furthermore, recent studies have already achieved groundbreaking results using machine learning to analyze MALDI-TOF MS data with microbes such as *Staphylococcus aureus*, reporting Area Under the Receiver Operating Characteristic (AUROC) scores up to 0.89 (Wang et al., 2022). AUROC is a commonly used metric in medicine to summarize the effectiveness of a model in predicting a binary outcome, which is calculated based on the area under the curve formed by the false positive rate and true positive rate as the model threshold is varied. Scores will always range from zero to one, corresponding to a model that incorrectly predicts every input and a model that correctly predicts every input.

Thus, to combat the emergence of *S. epidermidis* as a clinically relevant pathogen, a novel machine learning coupled with MALDI-TOF MS approach was taken to create prediction models for a variety of different antibiotics. Over 4,000 samples of *S. epidermidis* and their corresponding AMR profiles were taken from a mass spectra dataset compiled by Weis et al. (2022) from four different clinical sites between 2015 and 2018. In the study associated with the dataset, which contained data for multiple species, Weis et al. (2022) focused specifically on *Staphylococcus aureus*, *Escherichia coli*, and *Klebsiella pneumoniae* because it was found that species-specific predictions yielded high performance. However, they did not take *S. epidermidis* into account while developing and analyzing models for individual species. Furthermore, the priority pathogens evaluated in the study attained a maximum AUROC of 0.80 (Weis et al., 2022), which does not indicate poor performance but suggests room for improvement through additional measures.

Given the limitations of previous work and the lack of MALDI-TOF MS studies on nosocomial infections like *S. epidermidis*, the present study proposes a multitude of robust models for predicting the susceptibility of *S. epidermidis* to several antibiotics from various antibiotic families. SHapley Additive Explanations (SHAP) values were analyzed to improve model interpretations. The most significant feature bins for several models were examined and aligned with proteins listed in UniProt, confirming that the model’s selected features corresponded to combinations of relevant proteins associated with resistance. Results from the models exemplify not only a viable solution for improving existing methods for diagnosing and treating *S. epidermidis* infections but also a framework for developing MALDI-TOF MS diagnosis and prescription methodologies for other microbes in the future.

## 2. Materials and Methods

### 2.1 Source of Data

The present study utilizes an extensive mass spectra data collection of 303,195 microbes known as the Database of Resistance against Antimicrobials with MALDI-TOF Mass Spectrometry (DRIAMS) that was created by Weis et al. (2022) between 2015 and 2018. For each microbe analyzed, a corresponding instance exists in the DRIAMS dataset that contains the results of the MALDI-TOF MS analysis on the microbe and a laboratory-confirmed AMR profile. The dataset is also broken down into four sub-collections labeled DRIAMS-A to -D, each corresponding to a different site from which the data was collected. Of these sub-collections, DRIAMS-A has been identified to be the largest and most comprehensive subcollection, containing over 145,341 mass spectra in total. The clinical isolates presented in DRIAMS-A were collected by analyzing anonymized patient samples from the University Hospital Basel in Switzerland (Weis et al., 2022).

All data collection by Weis et al. (2022) followed standard ISO/IEC 17025 clinical procedures and was evaluated by the local ethical committee (IEC 2019-00729). The MALDI-TOF MS process was conducted using the Microflex Biotyper LT/SH System at the University Hospital of Basel, the site of the DRIAMS-A data collection. AMR profiles for several antibiotics (β-lactams [oxacillin, penicillin, amoxicillin-clavulanic acid, ceftriaxone], quinolones [ciprofloxacin], sulfonamides [cotrimoxazole], lincosamides [clindamycin], rifamycins [rifampicin], glycopeptides [teicoplanin], aminoglycosides [gentamicin], tetracyclines [tetracycline, tigecycline], and fusidanes [fusidic acid]) were constructed through traditional laboratory tests. After microbial identification and AMR profile compilation, each microbial instance was added to the corresponding DRIAMS dataset along with the mass spectra generated by MALDI-TOF MS (Weis et al., 2022). A processed version of the DRIAMS dataset was obtained from Kaggle with raw and preprocessed files that were unnecessary for model training removed (Scarlat, 2022). For each microbe sample, the downloaded dataset contains metadata, including the identified species, an AMR profile, and a sample ID (e.g., a0a359b5-c00a-4e6e-a195-728af31f7666) that corresponds to the file containing the mass spectra of the instance. For the present study, resistant and intermediate samples were labeled as positive classifications and susceptible samples as negative classifications.

### 2.2 Specifics of Mass Spectra Preprocessing in DRIAMS

The original dataset was preprocessed using mass spectra bins of 3 Da before machine learning analysis to balance peak separation and computational efficiency. Preprocessing was involved (1) intensity variance stabilization using the square root method, (2) smoothing with the Savitzky-Golay filter, (3) baseline removal with the statistics-sensitive non-linear iterative peak-clipping (SNIP) algorithm, (4) total ion current (TIC) calibration, (5) mass spectra trimming to a range of 2,000 to 20,000 Da. After that process, (6) each mass spectra was separated with a bin size of 3 Da, and (7) turned into a 6,000-dimensional vector ready for model training. Moreover, AMR resistance profiles for each antibiotic were initially reported by the assays to be either resistant, intermediate, or susceptible (Weis et al., 2022).

### 2.3 Workflow of the Present Study

The proposed workflow for the present study involves various processing, training, and interpretation steps (Figure 1). After obtaining the dataset, a random forest model was fitted to the data and combined with a meta-transformer to filter out irrelevant features. Six different classifiers were trained and tuned using filtered data, and then interpreted with biomarker analysis and external validation.

**Figure 1.**
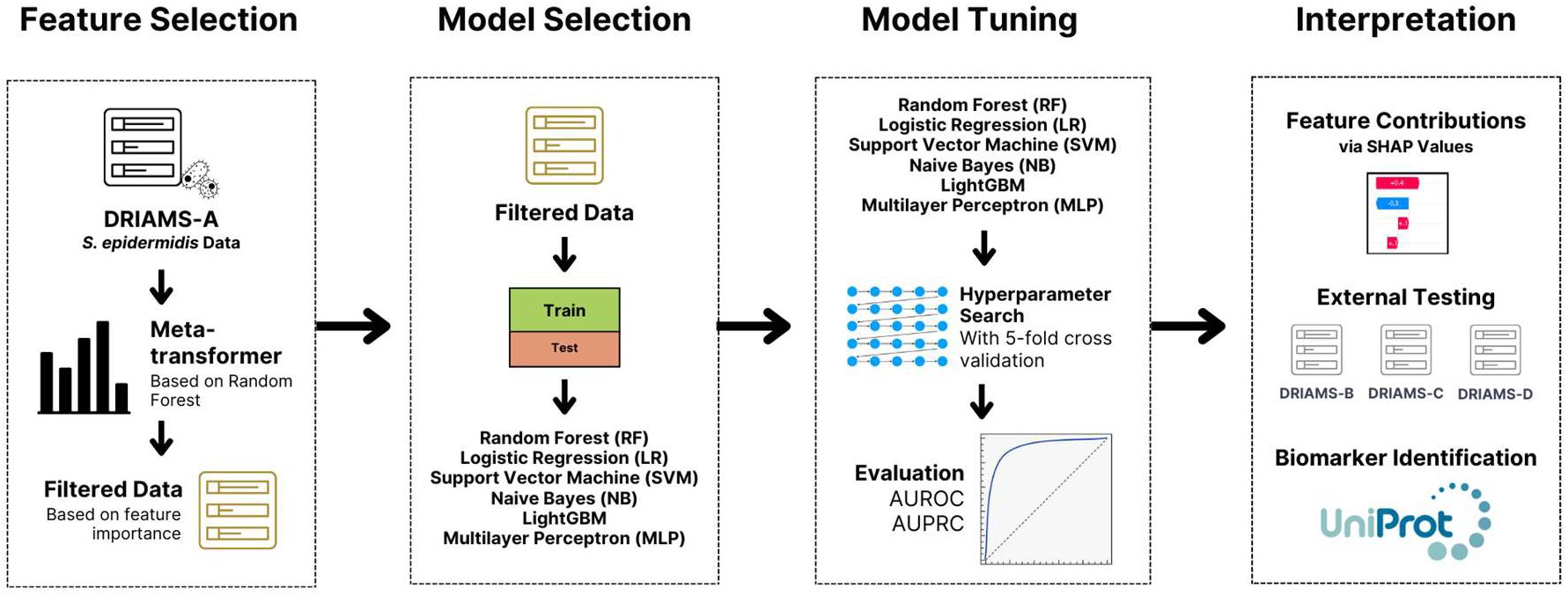
Summary of modeling workflow. *S. epidermidis* data is filtered with a meta-transformer and quickly tested with general, unoptimized models. This was followed by an extensive tuning process that aimed to optimize AUROC. After tuning, models were evaluated with SHAP values, which were subsequently matched with possible or known biomarkers. Finally, external testing on sites outside of DRIAMS-A was conducted.

### 2.4 Feature Selection

Datasets with too many irrelevant features can lead to redundant and overcomplicated algorithms with low accuracy and unnecessarily long training times. With over 6,000 mass spectra bins that profile the composition of each *S. epidermidis*, not every mass spectrum value generated by MALDI-TOF MS necessarily corresponds to a protein involved in imparting AMR to the bacterium. As such, feature selection becomes necessary to (1) reduce the number of considered features, (2) decrease the time required to train models, (3) reduce the likelihood of overfitting, and (4) avoid the curse of dimensionality that affects high-dimensional datasets like that of DRIAMS (Chen et al., 2020). A meta-transformer based on the Random Forest estimator was used to isolate relevant features and eliminate all other irrelevant ones (Feucherolles et al., 2022). The meta-transformer selects features by (1) fitting a Random Forest model to the entire dataset, (2) calculating the importance of each feature using the average reduction in impurity, (3) selecting relevant features based on a threshold with the 1.25 times the mean feature importance score as a baseline, and (4) reducing the dimensionality of the dataset by transforming the original dataset with irrelevant features removed.

### 2.5 Initial Model Selection

Each feature-selected mass spectra file was grouped into a single collection containing all mass spectra files for each unique antibiotic. These collections were subsequently split into a 70/30 training and test dataset with a stratification based on the susceptibility to the attributed antibiotic. For each antibiotic, models were constructed with the training datasets, while the test dataset was used for external validation on each trained model. A total of six different algorithms were implemented for each antibiotic collection: Random Forest (RF), Logistic Regression (LR), Naïve Bayes (NB), Support Vector Machine (SVM), LightGBM, and Multilayer Perceptron (MLP), as these models have performed well in previous machine learning studies on MALDI-TOF MS (Weis et al., 2022; Feucherolles et al., 2022). All classifiers except LightGBM were created using Scikit-learn with Python 3.11.3. Despite being installed as a separate library, LightGBM was able to integrate seamlessly with the Scikit-learn environment. Each of these models is robust and commonly used in machine learning for its ability to handle complex data, flexibility, and effectiveness in capturing complex relationships. RF is a popular ensemble learning technique that constructs numerous independent decision trees that are unique from one another. These trees are subsequently aggregated to generate predictions based on a given input. LR is a typical linear classifier that predicts the probability of an instance being positive or negative. Its linear nature makes it an excellent choice for accessible, efficient, and interpretable modeling of complex datasets. NB is akin to LR regarding ease of use and efficiency but builds on the assumption that the presence of one feature is unrelated to another (Feucherolles et al., 2022). SVM is known for its effectiveness in high-dimensional spaces, making it particularly useful for datasets with many features, such as mass spectra profiles. The linearity of data is not a significant concern because kernel tricks allow it to optimize hyperplanes that can separate complex relationships. Finally, LightGBM and MLP have been implemented in a large-scale MALDI-TOF MS study and performed the best out of all selected models (Weis et al., 2022). LightGBM is a gradient-boosting decision tree model that is not only fast and efficient but also scalable to large datasets. MLP is an artificial neural network that takes in data and passes it through hidden layers, allowing it to model complicated classification tasks accurately.

### 2.6 Model Tuning

After initial model selection, the best-performing model for each antibiotic was selected and optimized based on AUROC. AUROC was the evaluation metric of choice because it is not influenced by class ratio and provides an interpretable, overarching summary of performance with positive and negative susceptibility predictions for various classification thresholds. In addition, while many of the modeled antibiotics had relatively balanced datasets four antibiotic datasets had a positive class prevalence (PCP) of <0.20 or >0.80. Since AUROC is relatively invariant to the ratio of positive or negative classes, it is an acceptable metric for comparing the performance of different antibiotic models with different class ratios (Weis et al., 2022). Tuning was performed via hyperparameter optimization using Scikit-learn by exhaustively searching through a specified subset of the hyperparameter space—known as a grid—that is individually defined for each classifier. Validation of the model for each combination of hyperparameters in the grid was done using 5-fold cross-validation with a scoring method aiming to optimize AUROC. After the model tuning process, the Area Under the Precision-Recall Curve (AUPRC) metric was also used to evaluate the tuned models. AUPRC is another holistic metric used to assess model performance based on precision and recall across various thresholds and is especially useful for models trained on imbalanced datasets.

### 2.7 Finding and Interpreting SHAP Values

SHapley Additive exPlanations (SHAP) values can provide a more interpretable framework by quantifying the contribution of each feature to the classification outcome using a game theory approach (Shapley, 1953). Unlike Random Forest feature importances, which are usually based on the decrease in Gini impurity across all decision trees for each feature, SHAP values calculate the marginal contribution of each feature for every individual prediction. This can provide a more holistic view of all feature importances and illustrate the change in the expected model prediction for each specific feature. As such, SHAP values ensure fair interpretations in complex models where interactions between features can be non-linear and interdependent (Lundberg et al., 2017).

### 2.8 Biomarker Identification

Unlike decision trees, where the feature bins are ranked between 0 and 1, the range for possible SHAP values can vary widely depending on the model and data. However, higher absolute SHAP values always indicate a greater impact on model decisions. The most significant feature bins based on calculated SHAP values were investigated using UniProt and matched to the closest significant proteins around the range of Da values for each feature bin. This process helps identify novel resistance mechanisms and verify if the features identified by the model correspond to well-known and reputable mechanisms of antimicrobial resistance. UniProt analysis ensures that models are not simply capturing unclear patterns or noise within mass spectra profiles. In addition, it can also help identify connections with other related microbes like *S. aureus*.

### 2.9 External Validation

Models for each antibiotic are primarily created using DRIAMS-A because it is the most extensive and viable dataset out of DRIAMS- A-D. Given the independent construction of each DRIAMS dataset at distinct sites, this presents an ideal opportunity to evaluate the transferability of models across different clinical centers.

## 3. Results

### 3.1 Preprocessing and Feature Selection Summaries

After the *S. epidermidis* instances were preprocessed, a summary of all antibiotics of interest was obtained (Table 1). For 19 of the 20 antibiotics, the number of instances available for training and testing exceeded 4,000 samples. In the case of teicoplanin, only 488 samples were found in DRIAMS-A. The positive class prevalence (ratio of intermediate/resistant instances) was between 0.20 and 0.80 for 16 of the 20 antibiotics. However, the susceptibility status of most instances is heavily skewed towards one side for rifampicin and especially for ampicillin-amoxicillin, penicillin, and tigecycline. As such, it is necessary to avoid biased performance metrics such as accuracy and opt for metrics less sensitive to class distribution, such as AUROC and AUPRC. Another anomaly to consider with the summarized data is groups of antibiotics with an identical number of instances and positive class prevalence. In particular, one group of β-lactams that includes ceftriaxone all have a sample size of 4,261 and a positive class prevalence of 0.7055. In addition, both ampicillin-amoxicillin and penicillin have 4,371 instances and a positive class prevalence of 0.9938. Further analysis of DRIAMS-A indicates that for each instance that describes an antibiotic in either of the groups, the susceptibility of the instance to other antibiotics in the same group is identical. As such, only ceftriaxone and penicillin were considered to represent each of the respective groups because a model trained on any of the antibiotics within the same group would yield identical results.

**Table 1.**
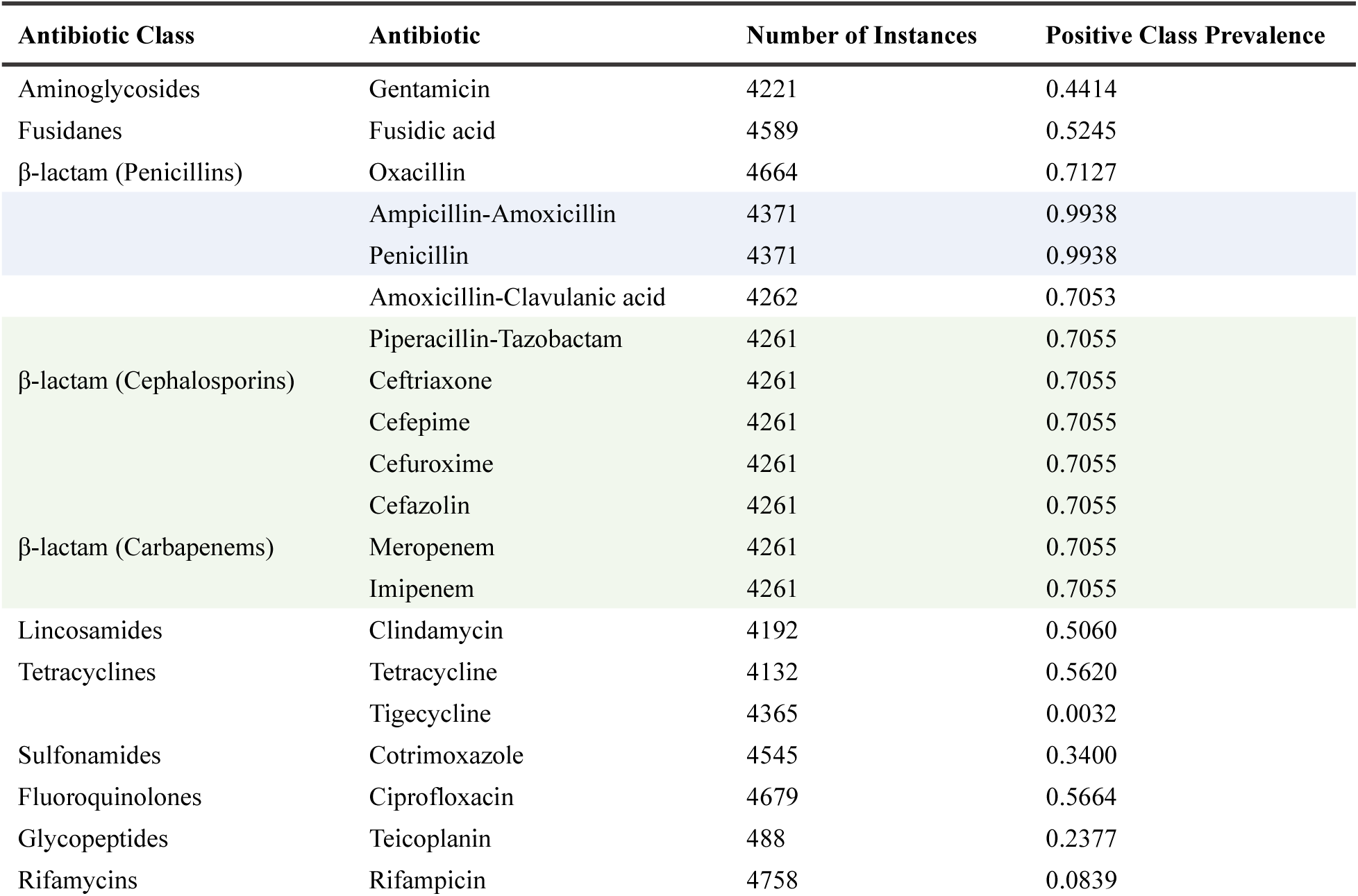
Summary of data for a variety of antibiotics evaluated in DRIAMS-A with sufficient sample size for creating admissible models to predict resistance in *S. epidermidis*. One group of β-lactam antibiotics is highlighted in light green because these antibiotics have the same number of instances, and each instance has the same resistance status for each antibiotic in the group. Another group of β-lactam antibiotics is highlighted in light blue for the same reasons.

After selecting one antibiotic from each identical instance group, a meta-transformer based on the Random Forest estimator was applied. The meta-transformer filtered out features with an information gain less than 1.25 times the mean. After feature selection, a summary of the antibiotic datasets (Table 2) indicates an average feature size of 1,158 across all antibiotic models after applying the meta-transformer.

**Table 2.** Summary of reduced model size after applying a Random Forest-based meta-transformer on the mass spectra. Originally a 6,000-dimensional vector, the number of features was reduced on average from 6,000 to 1,158. The highlighted rows indicate antibiotics selected to represent their respective groups from Table 1.

| Antibiotic | Original Feature Size | Number of Selected Features |
| --- | --- | --- |
| Gentamicin | 6000 | 989 |
| Fusidic acid | 6000 | 1375 |
| Oxacillin | 6000 | 1182 |
| Penicillin | 6000 | 1324 |
| Amoxicillin-Clavulanic acid | 6000 | 1206 |
| Ceftriaxone | 6000 | 1195 |
| Clindamycin | 6000 | 1376 |
| Tetracycline | 6000 | 489 |
| Tigecycline | 6000 | 923 |
| Cotrimoxazole | 6000 | 956 |
| Ciprofloxacin | 6000 | 848 |
| Teicoplanin | 6000 | 1645 |
| Rifampicin | 6000 | 1545 |

### 3.2 Model Creation

Six different machine learning models were initially assessed for each antibiotic in this study, including Random Forest (RF), Logistic Regression (LR), Naïve Bayes (NB), Support Vector Machine (SVM), LightGBM, and Multilayer Perceptron (MLP). Initially, the datasets for *S. epidermidis* were stratified by susceptibility and randomly partitioned into a 70/30 split for training and testing, respectively, to maintain the integrity of class proportions. Default hyperparameters were applied to train each model on training data, while testing data was used to evaluate the models’ effectiveness on microbial instances the model had not encountered. For the most part, the best-performing models were RF, SVM, and LightGBM when considered across every collection of antibiotic data (Table 3). Interestingly, MLP was not able to keep up with these classifiers, despite being a neural network model. However, such a summary can only be used as a reference, as the best-performing model can vary and necessitates individual tuning for each antibiotic data collection.

**Table 3.** Summary of default hyperparameters that were tuned later in the study. Average AUROC scores associated with the default hyperparameters for each classifier across all evaluated antibiotic data collections are also reported.

| Classifier | Default Hyperparameters | Average AUROC Score |
| --- | --- | --- |
| Random Forest (RF) | n_estimators=100<br>max_features="sqrt" | 0.81 |
| Logistic Regression (LR) | C=1.0<br>penalty="L2"<br>solver="lbfgs"<br>max_iter=100 | 0.69 |
| Support Vector Machine (SVM) | C=1.0<br>gamma="scale"<br>kernel="rbf" | 0.84 |
| Naïve Bayes (NB) | - | 0.71 |
| LightGBM | boosting_type="gdbt"<br>n_estimators=100<br>learning_rate=0.1 | 0.82 |
| Multilayer Perceptron (MLP) | hidden_layer_sizes=(100,)<br>max_iter=200 | 0.73 |

After initial model evaluation, a multi-faceted model tuning process was carefully conducted, extensively searching a variety of hyperparameter combinations to improve model performance. A summary of tuned hyperparameters is available (Supplemental Figure 1). All six models mentioned above were tuned for each antibiotic data collection using 5-fold cross-validation based on finding a more optimized AUROC score. Tuned hyperparameters for each classifier are listed in the previous table. Generally, LightGBM and SVM were the best-performing classifiers, but other models also performed relatively well (Figure 2). Also, models maintained high AUPRC scores despite being trained to optimize AUROC. This indicates that they maintained precision and recall balance across various thresholds, reflecting a consistent ability to distinguish between the classes even with imbalanced datasets. After comparing the performance of different classifiers for the three selected antibiotics, all antibiotic models were fitted to the best-performing classifier along with the tuned hyperparameters. All models performed exceptionally well, achieving mean AUROC scores ranging from 0.80 to 0.95 after ten separate, shuffled, and stratified 70/30 train-test splits were performed on each (Figure 3). The best predictive performance was achieved by either LightGBM (7 models) or SVM (6 models) for all antibiotic data collections. Additionally, the standard deviations were generally low between each split except for penicillin and tigecycline. However, it is worth noting that the PCP for their respective data collections was 0.9938 and 0.0032, which accounts for the high variation in model performance between each test because there is always very little of either the positive or negative class to train on.

**Figure 2.**
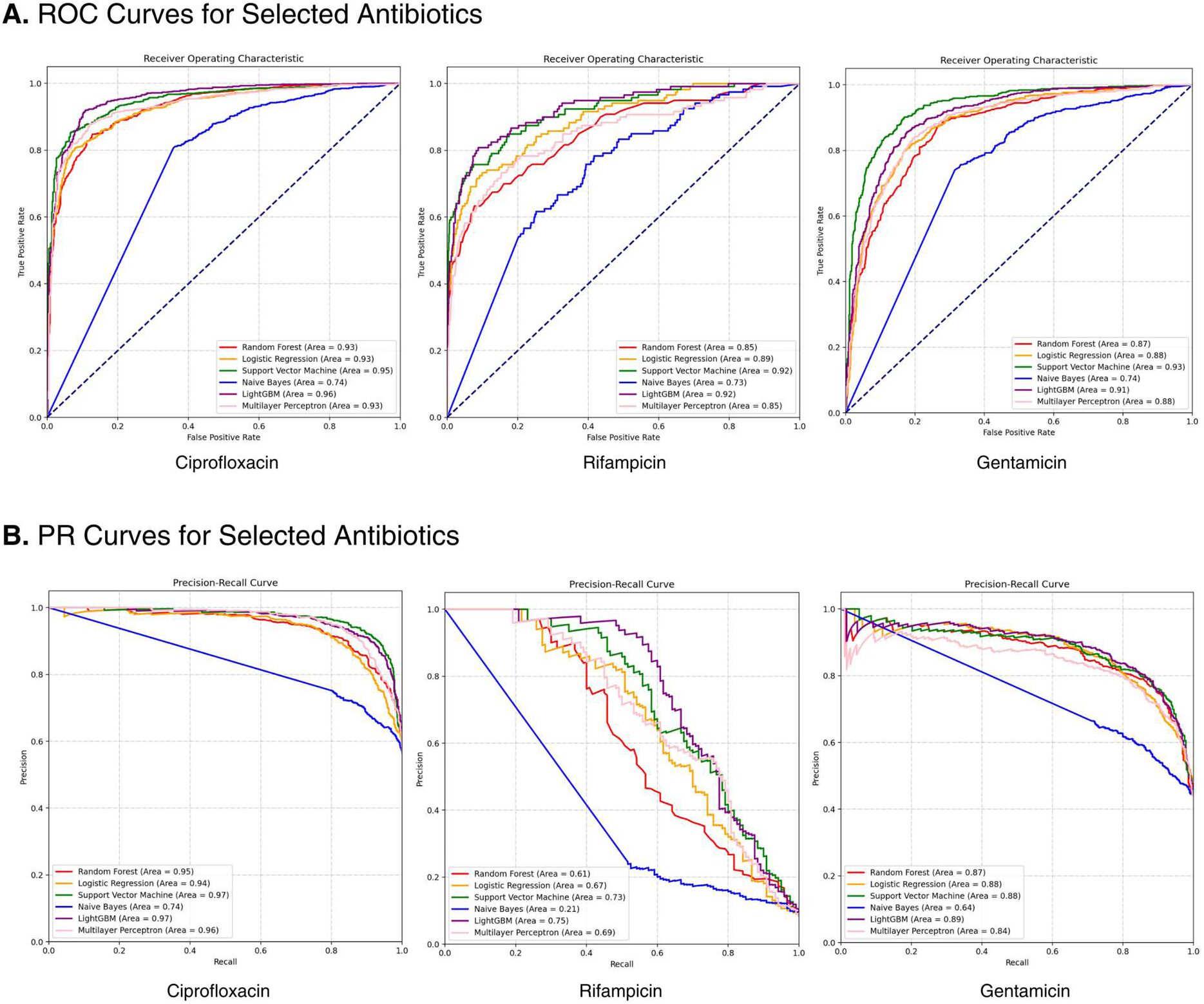
Performance of the six classifiers was evaluated for three selected antibiotics—ciprofloxacin, rifampicin, and gentamicin—and yielded high performance for both AUROC and AUPRC metrics in a single 70/30 train-test split. The graphs represent the performance of tuned models in predicting *S. epidermidis* resistance to the drug on each respective antibiotic data collection. **A. Comparison of receiving operating characteristic curves for models under varying thresholds.** For ciprofloxacin, LightGBM had the highest predictive performance (AUROC = 0.96). On the other hand, both LightGBM and SVM performed equally well for rifampicin (AUROC = 0.92), while SVM alone performed the best for gentamicin (AUROC = 0.93). **B. Comparison of precision-recall curves for models under varying thresholds.** Unlike AUROC, which is evaluated using 0.5 as the baseline, AUPRC is evaluated using the positive class prevalence (PCP) as the baseline. LightGBM performed the best for all three selected antibiotics for AUPRC. Ciprofloxacin (PCP = 0.57) was able to achieve high predictive performance (AUPRC = 0.97). The overall performance for rifampicin (PCP = 0.08) was also extremely high (AUPRC = 0.75) despite being noticeably lower than ciprofloxacin because of rifampicin’s low PCP baseline. Gentamicin (PCP = 0.44) also performed relatively well using AUPRC as a metric (AUPRC = 0.89).

**Figure 3.**
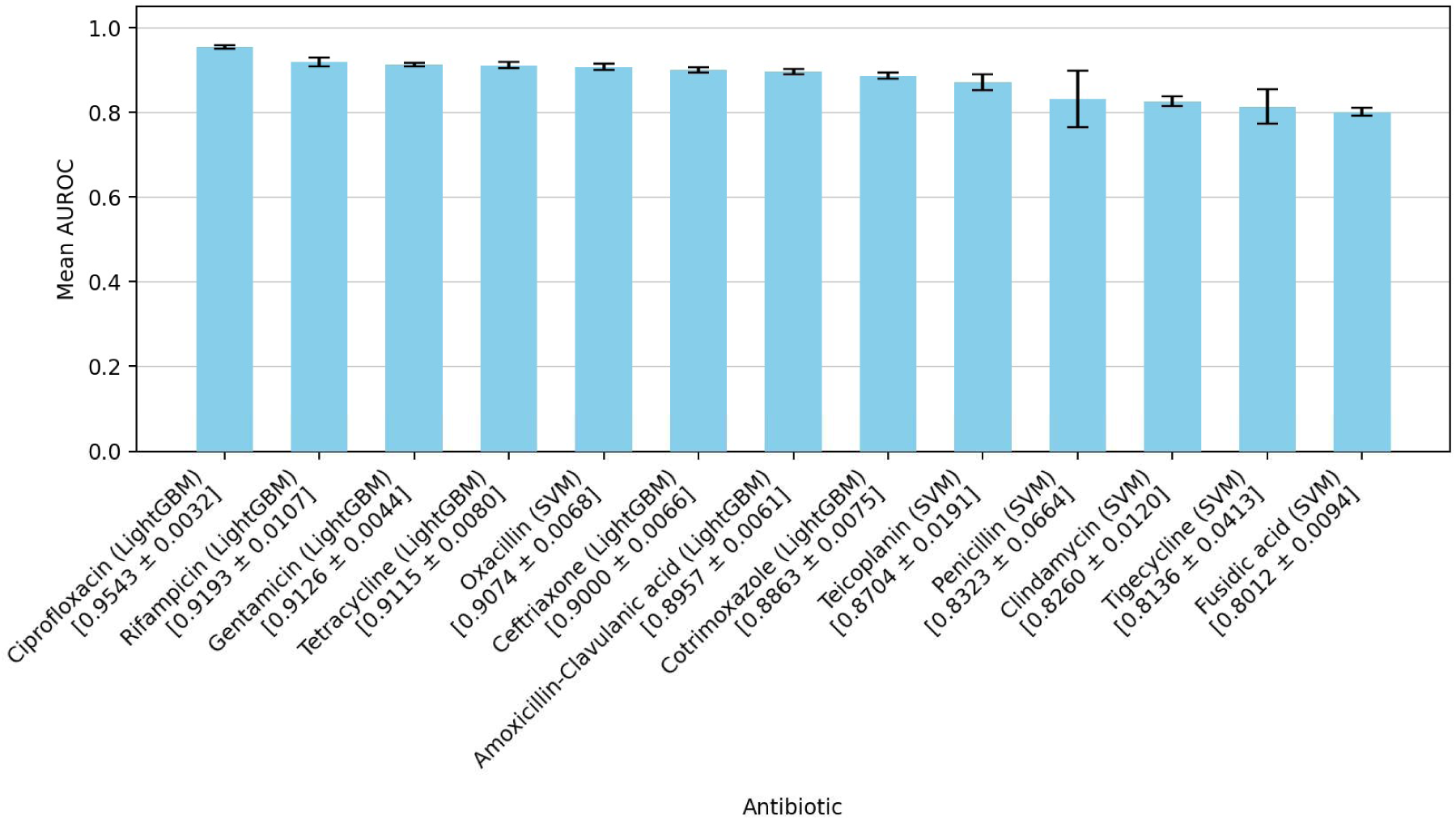
Average performance based on AUROC for machine learning models across all antibiotic data collections. Bars report the mean ± standard deviation for ten separate, shuffled, and stratified 70/30 train-test splits.

The PCP for a collection of samples had a significant impact on AUPRC as well. Although ROC curves were similar for the best-performing classifiers, there were some noticeable differences in the PR curves for the classifiers that performed well with respect to AUROC (Figure 4). For example, although the best-performing model for rifampicin achieved a high AUROC of 0.94, the same model attained an AUPRC of 0.80. Compared to gentamicin, whose model had a similar AUROC of 0.93, an AUPRC of 0.80 is noticeably lower than gentamicin’s AUPRC of 0.91. Despite this, the performance of rifampicin can still be justified due to the differing baselines for the two models. In fact, unlike AUROC, which uses a baseline of 0.5, the baseline for AUPRC is the PCP of the corresponding dataset. Because the PCP of rifampicin is low at 0.08, an AUPRC of 0.80 is commendable and indicates adequate model performance.

**Figure 4.**
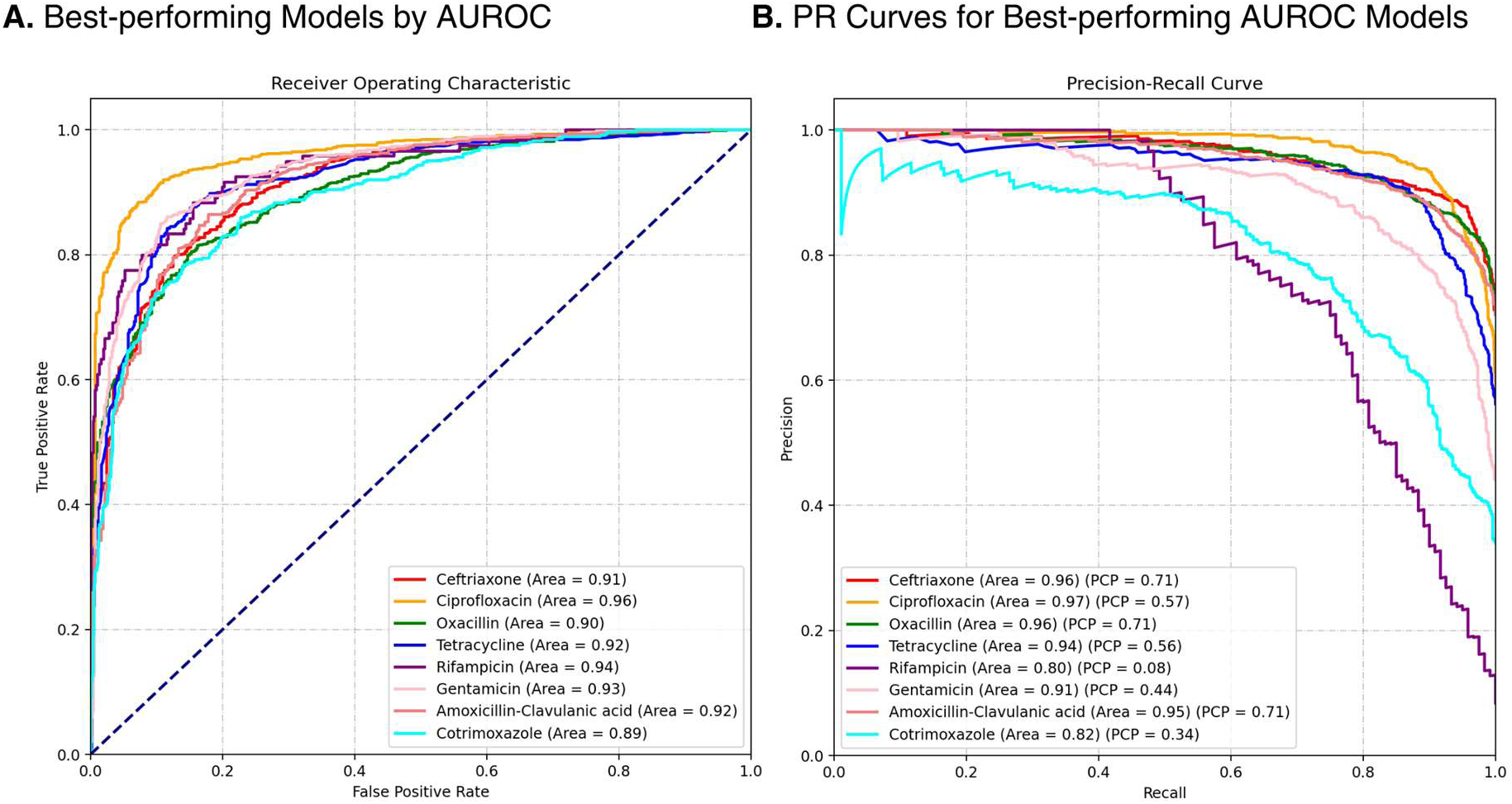
Performance of eight different antibiotics plotted against one another under the ROC and PR curves. **A. ROC curves and corresponding areas for the best-performing model for each antibiotic data collection.** The model was selected by finding the train-test split with the highest AUROC performance across selected antibiotics. **B. Corresponding AUPRC for the same models selected previously.** Positive class prevalence (PCP) is also listed to serve as a baseline for comparing model performances based on AUPRC.

Among the selected antibiotics, ciprofloxacin, rifampicin, and gentamicin performed the best, with AUROC scores ranging from 0.93 to 0.96. Other antibiotic models including the models for tetracycline, amoxicillin-clavulanic acid, ceftriaxone, and oxacillin also performed well, maintaining AUROC scores of at least 0.90. On the other hand, ciprofloxacin, ceftriaxone, and oxacillin performed the best for AUPRC, with scores ranging from 0.96 to 0.97. The model for ciprofloxacin was the most effective for both metrics, with a maximal AUROC of 0.96 and AUPRC of 0.97. Rifampicin was not far behind, achieving a maximal AUROC of 0.94 but an AUPRC of 0.80.

Moreover, despite being the least performative models on average, the models constructed for fusidic acid, tigecycline, and clindamycin maintained an average AUROC between 0.80 and 0.83, which is considered acceptable performance under most circumstances. Overall, the performance of such a multitude of antibiotics, when analyzed with machine learning, illustrates the potential for MALDI-TOF MS to emerge as a leading technique in diagnosing bacterial infections like *S. epidermidis*. A summary of all performance metrics is also available (Supplemental Figure 2).

### 3.3 Interpretation of SHAP Values and Biomarker Identification

Evaluation of feature importance based on SHAP was conducted on the best-performing models of ciprofloxacin, gentamicin, and rifampicin. SHAP values help explain the relevance of a feature bin and provide a deeper understanding of the biological mechanisms behind antibiotic resistance in S. epidermidis. In the case of the ciprofloxacin, the feature bins of 1,916 and 1,604 appear to have contributed the most to the output of the model (Figure 5). A pseudogel plot of a randomly selected susceptible ciprofloxacin instance and resistant ciprofloxacin instance illustrates the locations of these two feature bins, where there is a noticeable difference in the intensity of protein peaks at each feature bin can be observed (Figure 6). As such, the approximate location of these feature bins likely contains a biomarker associated with the impartment of antibiotic resistance. Closely associated proteins to these feature bins have been identified around mass values of 7,753 Da and 6,812 Da. The same process was used to analyze the five most important feature bins for gentamicin and rifampicin, with associated protein names and UniProt IDs recorded (Table 4). Out of the fifteen feature bins, five of them did not contain a documented S. epidermidis protein within the range, so a closely associated protein in S. aureus was instead chosen. In addition, six out of the fifteen feature bins corresponded to an uncharacterized protein in S. epidermidis, indicating the possibility for further research into these biomarkers.

**Figure 5.**
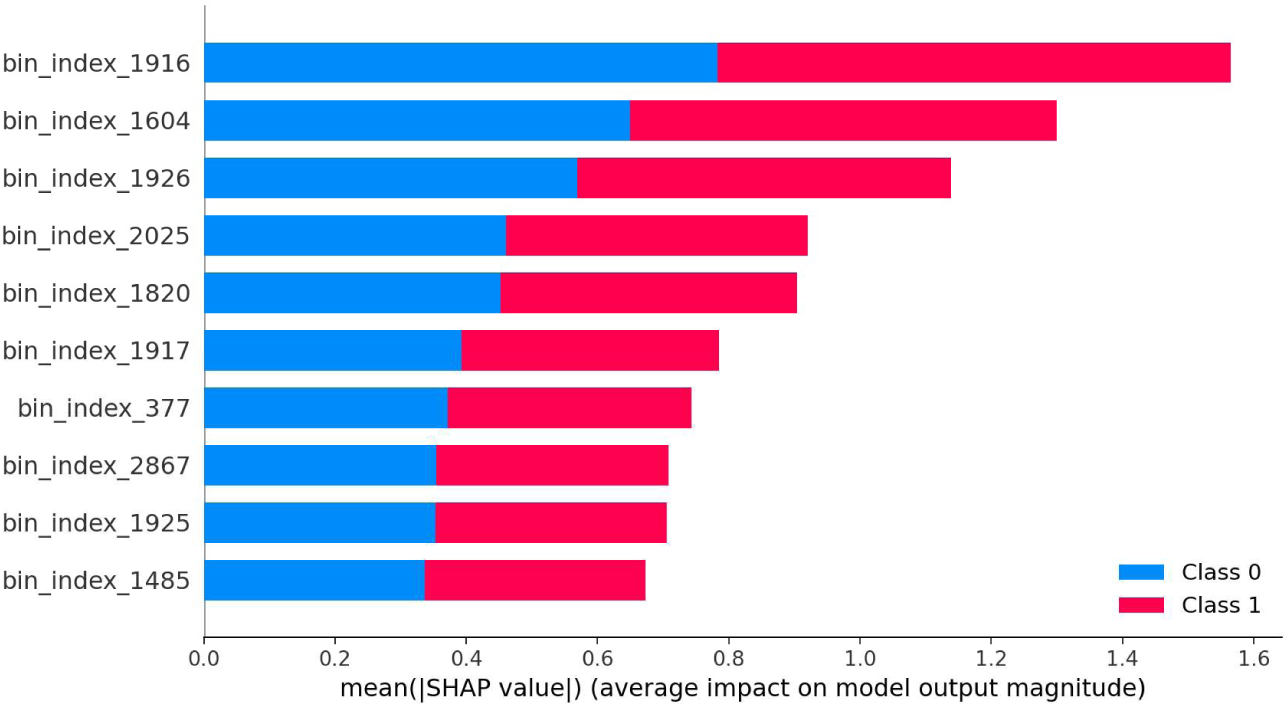
List of most important features by contribution to ciprofloxacin model based on SHAP values. Longer bars generated by the program indicate that a feature has a more considerable influence on model output. Most features appear to be located around feature bin 2,000, which corresponds to a Da value of approximately 8,000.

**Figure 6.**
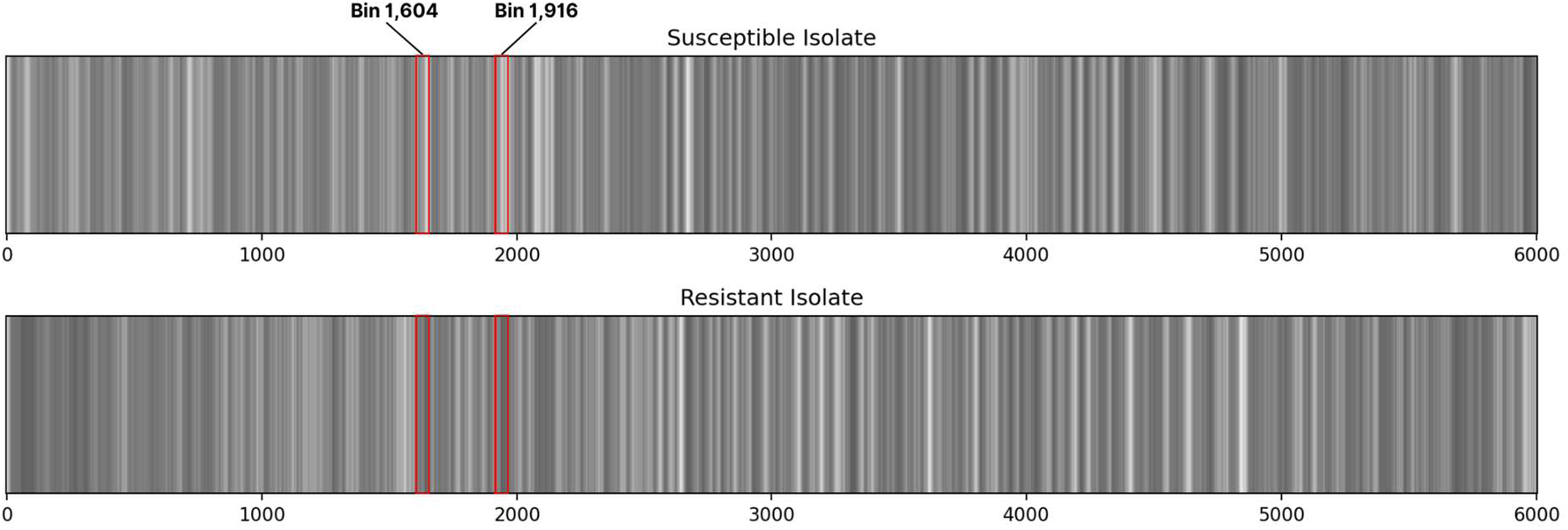
Pseudogel plot of two randomly selected ciprofloxacin isolates of different classes from DRIAMS-A. The approximate locations for two bins are shown. For bins 1,916 and 1,604, there appears to be a peak (darker intensity) in the resistant isolate but not the susceptible, indicating the existence of a significant protein biomarker for antibiotic resistance.

**Table 4.** Most important feature bins and closest significant protein biomarkers for the best-performing models. The closest significant protein was found by (1) searching for the best-annotated *S. epidermidis* protein within the feature bin and (2) identifying the best-annotated protein within the feature bin for *S. aureus*, a closely related bacteria if an entry could not be found for *S. epidermidis*. Listed masses and annotations were acquired directly from the UniProt entry for each protein. Any proteins identified from *S. aureus* are marked with an asterisk* in the UniProt ID column.

| Antibiotic | Rank | Feature Bin | Mass (Da) | UniProt Annotation | UniProt ID |
| --- | --- | --- | --- | --- | --- |
| Ciprofloxacin | 1 | 1,916 | 7,748 | Uncharacterized protein (M453_0201920) | A0A829M6R5 |
|  | 2 | 1,604 | 6,812 | UPF0337 protein SE_0604 | Q8CTA8 |
|  | 3 | 1,926 | 7,780 | Uncharacterized protein (I3V53_12125) | A0A8I1BC98 |
|  | 4 | 2,025 | 8,077 | Pathogenicity island family protein | A0A0E8JV84* |
|  | 5 | 1,820 | 7,460 | Uncharacterized protein | A0A125SJI8 |
| Gentamicin | 1 | 1,917 | 7,753 | Transposase | A0A2G7I0T7 |
|  | 2 | 2,411 | 9,233 | RecG wedge domain-containing protein | A0A0E0VNH2* |
|  | 3 | 326 | 2,979 | Delta-hemolysin | P0C1V1* |
|  | 4 | 2,857 | 10,572 | LPXTG-motif cell wall anchor domain protein | A0A0E1VM26* |
|  | 5 | 625 | 3,877 | Uncharacterized protein (SERP0759) | Q5HPZ8 |
| Rifampicin | 1 | 2,025 | 8,077 | Pathogenicity island family protein | A0A0E8JV84* |
|  | 2 | 2,035 | 8,107 | Pathogenicity island protein | G9D5B0 |
|  | 3 | 1,242 | 5,726 | Uncharacterized protein (I3V53_09740) | A0A6B9V0A1 |
|  | 4 | 4,736 | 16,210 | Uncharacterized protein (I3V53_10930) | A0A8I0WAC7 |
|  | 5 | 1,922 | 7,766 | DNA-directed RNA polymerase subunit omega | Q5HPY0 |

### 3.4 External Validation

After tuned models for each antibiotic were optimized, isolates from DRIAMS- B-D were tested on models trained on their entire corresponding DRIAMS-A dataset. Overall, performance varied greatly from site to site, with some models carrying over their performance between sites and others performing less optimally (Table 5). Other times, there was not sufficient or workable data for *S. epidermidis* in other sites to analyze the performance of different antibiotic models properly.

**Table 5.** AUROC score for best-performing models of each antibiotic in DRIAMS-A when evaluated using an external site. The number of samples available for testing is given in the title of each column. When the prevalence of the positive class is either 0 or 1, or if there is no data available on the antibiotic in the dataset, nothing is recorded because calculating AUROC would not be meaningful. The best-performing entries based on AUROC are highlighted in light green.

| Antibiotic | DRIAMS-B (n=213) | DRIAMS-C (n=21) | DRIAMS-D (n=374) |
| --- | --- | --- | --- |
| Gentamicin | 0.452 | 0.702 | 0.704 |
| Fusidic acid | 0.557 | 0.250 | - |
| Oxacillin | 0.374 | 0.613 | - |
| Penicillin | - | 0.150 | 0.442 |
| Amoxicillin-Clavulanic acid | 0.353 | 0.963 | - |
| Ceftriaxone | - | - | - |
| Clindamycin | 0.328 | 1.000 | 0.605 |
| Tetracycline | 0.306 | - | 0.813 |
| Tigecycline | - | - | 0.946 |
| Cotrimoxazole | - | 0.683 | 0.748 |
| Ciprofloxacin | 0.598 | 1.000 | 0.879 |
| Teicoplanin | - | - | - |
| Rifampicin | 0.776 | - | 0.717 |

## 4. Discussion

Many studies have shown that MALDI-TOF MS is not only a faster approach than existing AST methods but also much more cost-effective (Dhiman et al., 2011). In order to leverage the full capabilities of MALDI-TOF MS in AST, combining it with other analysis techniques, such as machine learning after dimensional reduction, is necessary to extract meaningful information for resistance predictions. With high-performing models, such a workflow would be relevant and reasonably surpass traditional AST methods in practice. Although several other studies on machine learning applications in MALDI-TOF MS have already been conducted on various pathogens, there are no studies at the time of writing that specifically address nosocomial infections such as *S. epidermidis*. Furthermore, most publications focus only on one antimicrobial class at a time (e.g., glycopeptides like vancomycin) rather than a more diverse group of differing antimicrobials necessary for a holistic evaluation (Feucherolles et al., 2022). This reduces the impact of such research on antibiotic stewardship because these strong antibiotics are usually last-resort choices for patients unable to recover using more common antibiotics due to the prevalence of multidrug-resistant bacteria. Frequent and indiscriminate use of strong antibiotics like vancomycin increases the prevalence of resistant bacteria, so using these drugs only when necessary helps slow the emergence of superbugs (Remschmidt et al., 2017). The effectiveness of these last-resort drugs is illustrated with *S. epidermidis* in DRIAMS-A as well—nearly all isolates were susceptible to vancomycin, so model creation would not necessarily be advantageous.

As such, this study serves as a consideration of the potential for mass spectrometry to be combined with machine learning to rapidly predict AMR for a variety of different antibiotics in nosocomial infections. The high AUROC scores up to 0.95 and AUPRC scores up to 0.97 achieved by this study can be attributed either to the quality and scale of the *S. epidermidis* dataset or the usage of feature selection as a technique to eliminate irrelevant features and prevent the creation of an overcomplicated model. Indeed, the original study conducted by the authors of the DRIAMS-A dataset achieved maximal AUROC scores of 0.74, 0.74, and 0.80 for *Staphylococcus aureus*, *Escherichia coli*, and *Klebsiella pneumoniae*, respectively (Weis et al., 2022), which are all lower than or equal to the lowest average AUROC score of 0.80 for fusidic acid in the antibiotics modeled for *S. epidermidis.* In addition, their corresponding AUPRC scores were 0.49, 0.30, and 0.33, respectively, which are significantly overshadowed by the near-perfect AUPRC scores achieved by the best-performing *S. epidermidis* models. Although the results achieved by this study were significantly more performative than those achieved in the original study, it is essential to remember that the microbe of focus is different in this study, and other factors may have been involved in the performance of these models.

Another detail to note is that Weis et al. (2022) did not mention feature selection before constructing models for each antibiotic, which could have reduced the efficiency of their models. RF was specifically chosen for feature selection due to its advantages over other methods. For one, previous studies have shown that the performance of models trained with features filtered by RF has outperformed those filtered by other classifiers like K-Nearest Neighbors (KNN), Linear Discriminant Analysis (LDA), and SVM (Chen et al., 2020). In addition, although SHAP was later used for model explanation, it was not used for feature selection because the time required for SHAP analysis increases significantly with each added column—by almost 80%—making it unsuitable for extremely high-dimensional datasets (Ebrahimi and Patel, 2022). RF already provides sufficient performance for preliminary feature filtering and reduces the number of features to a size that can be more efficiently handled by SHAP analysis.

As shown in Figure 3, despite high average AUROC scores, the classifiers for penicillin and tigecycline had relatively high standard deviations for AUROC scores between the ten separate train-test splits. Such performance is likely explained by the fact that these classifiers are subject to highly imbalanced class prevalences, with both approaching nearly no susceptible or resistant isolates, respectively. Out of the 4,371 instances in DRIAMS-A, only 27 isolates of *S. epidermidis* were susceptible to penicillin. Similarly, out of the 4,365 instances of tigecycline in DRIAMS-A, only 14 isolates were resistant to the tigecycline. In many cases, imbalanced datasets can be fixed using oversampling techniques such as Synthetic Minority Oversampling Technique (SMOTE). However, due to the high dimensionality of each mass spectra instance, consequential data points are often sparsely distributed, making it difficult for SMOTE to generate meaningful synthetic samples and improve model performance. Past studies with other high dimensional datasets like genome data have demonstrated this, showing negligible improvement in model performance after applying SMOTE (Blagus and Lusa, 2012). On the other hand, undersampling techniques can lead to significant information loss. Due to the overwhelming difference in numbers between the positive and negative classes, discarding majority class instances to match a small minority can result in a significant loss of diversity, crucial insights, and meaningful patterns, with these effects being exacerbated by the gap in the number of samples in each class. As such, holistic metrics like AUROC and AUPRC are crucial for capturing and evaluating model performance based on both the majority and minority classes at the same time.

SHAP analysis provided insight into novel, uncharacterized biomarkers as well as evidence for the functions of proteins that have already been documented. For example, the feature bin of 1,916 in the ciprofloxacin model contributed significantly to model performance and was identified to be a known but minimally documented protein at 6,812 Da (Uncharacterized protein (M453_0201920)). Moreover, the second most relevant feature for predicting ciprofloxacin resistance was calculated to be the feature bin of 1,604, which corresponds to UPF0337 protein SE_0604 at 6,812 Da. Belonging to the UPF0337 (CsbD) family, a MALDI-TOF MS different study conducted on *S. aureus* found that another UPF0337 protein that underwent a mutation reduced antibiotic binding affinity through a molecular docking simulation, confirming that mutations may also play a major role in antibiotic resistance (Yu et al., 2022). On the other hand, another possibility that explains the significance of particular feature bins is the horizontal transfer of resistance-encoding genes between Staphylococci or other bacterial species, such as the closely related *S. aureus*. For example, another relevant feature for the ciprofloxacin model corresponding to the feature bin of 2,025 was not identified for *S. epidermidis* in UniProt but was identified at 8,077 Da as a pathogenicity island family protein in *S. aureus*. Pathogenicity island proteins are known to cause virulence and impart antibiotic resistance to microbes that were not initially resistant and have been documented to be transferred by other studies (Bloemendaal et al., 2010). In the case of gentamicin, the most relevant feature bin of 1,917 was identified as transposase at 7,753 Da. Transposase is responsible for the movement of transposons—which may encode antibiotic resistance—from one plasmid to another (Babakhani and Oloomi, 2018). Another interesting feature identified by SHAP analysis was the feature bin of 326, for which, although no documented protein exists for *S. epidermidis* in UniProt, it corresponds to a delta-hemolysin biomarker at 2,979 Da in *S. aureus*. Delta-hemolysin is one of the primary virulence agents of *S. aureus*, is known to increase bacterial resistance to antibiotics indirectly, and has also been identified in some isolates of *S. epidermidis* (Motamedi et al, 2018; Pinheiro et al., 2015). This illustrates the significance of gene transfers between Staphylococci in AMR while also emphasizing the ability of MALDI-TOF MS machine learning models to capture relevant biomarkers rather than noise. Finally, the same pathogenicity island family protein identified by SHAP analysis in ciprofloxacin was also the most relevant feature of the rifampicin model, demonstrating that AMR resistance is multifaceted and similar resistance mechanisms exist between different families of antibiotics. Additionally, the rifampicin model also detected the presence of DNA-directed RNA polymerase subunit omega—encoded by the *rpoZ* gene—at 7,766 Da for the feature bin of 1,922. Recently, the *rpoZ* gene and the subunit encoded by it were validated to be a significant factor in biofilm formation, motility, and antibiotic resistance, although in *E. coli* and not *S. epidermidis* (Patel, 2020).

However, interpretation of the roles of the features within the bacteria should be made cautiously due to limitations. Due to the restrictions of MALDI-TOF MS, it is more challenging to detect proteins with larger masses. The current dataset is limited to 20,000 Da, meaning that instead of detecting the true protein involved in resistance, the models likely detect a different protein or peptide encoded within the same plasmid involved in the same antibiotic resistance mechanism (Lau et al., 2014). For example, previous studies have demonstrated that the primary mutation responsible for ciprofloxacin resistance are mutations in the *gyrA* and *parC* genes, which have masses of 100,144 Da and 91,145 Da, respectively (Kang et al., 2020). Such proteins are outside of the detection range and are therefore not considered when building the model, demonstrating that although model performance may not be optimized to detect the main resistance biomarker directly, highly accurate results can still be achieved by detecting other smaller but related proteins.

There are several other limitations in the present study. First, dealing with imbalanced datasets is challenging due to the dimensionality of mass spectra. As such, constructing models to analyze MALDI-TOF MS may not be adequate when an antibiotic is nearly always resisted or effective against *S. epidermidis*. In addition, models trained on one site may not easily transfer to another—even though processes can be nearly identical between sites. There are often other factors, such as differing calibrations of the same instruments, that lead to compounding differences between the mass spectra generated at one site compared to the mass spectra generated at another. This can be seen in the ciprofloxacin resistance model, where the model trained on DRIAMS-A performed relatively well on DRIAMS-C and DRIAMS-D but was not as adequate in DRIAMS-B.

While these limitations are known and should be considered with care, the economic and human benefits of a successful implementation are widespread. For example, in a retrospective clinical case study, Weis et al. (2022) provided treatment guidance up to 72 hours earlier than traditional AST approaches, significantly impacting treatment workflow through AMR predictions based on MALDI-TOF MS. In fact, 88% of the time, using AMR predictions generated by machine learning models benefitted the patient and promoted antibiotic stewardship (Weis et al., 2022). Rather than completely replacing traditional AST methods, MALDI-TOF MS should be used to give early guidance in drug prescription until confirmatory results from traditional AST are attained (Feucherolles et al., 2022).

## 5. Conclusion

When combined with machine learning, MALDI-TOF MS has proven to be a powerful tool for rapid, economical, and efficient AMR prediction in *S. epidermidis* for a variety of different antibiotics and antimicrobial families. With the amount of predictive power that can be derived from a single MALDI-TOF MS diagnostic for so many antibiotics, MALDI-TOF MS has the potential to serve as a holistic, go-to tool for species identification, AMR profile prediction, and diagnostic for identifying novel biomarkers for future investigation. As long as sufficient data is available with ample representation of both the susceptible and resistant isolates, MALDI-TOF MS can be combined with machine learning to benefit the patient-clinician workflow and contribute to antibiotic stewardship efforts. This study serves as a significant example for future research to follow, which should aim to investigate a broader range of species, increase the mass spectra range, integrate patient-specific data such as family history to create an all-encompassing model, and investigate potential biomarkers identified by the machine learning models.

