## Supplemental Figure for "Elucidating Resistance Mechanisms in *Staphylococcus epidermidis*: A High-Performing MALDI-TOF MS-Based Proteomic Approach for Predictive Modeling"

### Supplemental Material

The hyperparameters for the corresponding classifiers are summarized (Supplemental Figure 1). Another summary of the performance of all classifiers and the exact mean and standard deviations for AUROC and AUPRC was compiled (Supplemental Figure 2).

**Supplemental Figure 1 |** Tuned hyperparameters of best-performing classifiers after an extensive grid search. Each set of hyperparameters corresponds to the same antibiotic classifier in Supplemental Figure 1.

| Antibiotic | Tuned Hyperparameters |
|---|---|
| Gentamicin | boosting_type: "gbdt", learning_rate: 0.1, n_estimators: 400, objective: "binary" |
| Fusidic acid | C: 1, gamma: "auto", kernel: "rbf", objective: "binary", probability: True |
| Oxacillin | C: 10, gamma: "auto", kernel: "rbf", objective: "binary", probability: True |
| Penicillin | C: 10, gamma: "scale", kernel: "rbf", objective: "binary", probability: True |
| Amoxicillin-Clavulanic acid | boosting_type: "gbdt", learning_rate: 0.1, n_estimators: 400, objective: "binary" |
| Ceftriaxone | boosting_type: "gbdt", learning_rate: 0.1, n_estimators: 400, objective: "binary" |
| Clindamycin | C: 1, gamma: "auto", kernel: "rbf", objective: "binary", probability: True |
| Tetracycline | boosting_type: "gbdt", learning_rate: 0.1, n_estimators: 100, objective: "binary" |
| Tigecycline | C: 10, gamma: "scale", kernel: "rbf", objective: "binary", probability: True |
| Cotrimoxazole | boosting_type: "gbdt", learning_rate: 0.1, n_estimators: 400, objective: "binary" |
| Ciprofloxacin | boosting_type: "gbdt", learning_rate: 0.1, n_estimators: 400, objective: "binary" |
| Teicoplanin | C: 10, gamma: "scale", kernel: "rbf", objective: "binary", probability: True |
| Rifampicin | boosting_type: "gbdt", learning_rate: 0.1, n_estimators: 400, objective: "binary" |

**Supplemental Figure 2 |** Summary of all antibiotic model classifiers, mean AUROC score, and mean AUPRC score after ten separate, stratified, and shuffled 70/30 train-test splits.

| Antibiotic | Classifier | AUROC | AUPRC |
|---|---|---|---|
| Gentamicin | LightGBM | 0.9126 ± 0.0044 | 0.8783 ± 0.0111 |
| Fusidic acid | Support Vector Machine | 0.8012 ± 0.0094 | 0.8018 ± 0.0108 |
| Oxacillin | Support Vector Machine | 0.9074 ± 0.0068 | 0.9543 ± 0.0056 |
| Penicillin | Support Vector Machine | 0.8323 ± 0.0664 | 0.9990 ± 0.0006 |
| Amoxicillin-Clavulanic acid | LightGBM | 0.8957 ± 0.0061 | 0.9495 ± 0.0077 |
| Ceftriaxone | LightGBM | 0.8998 ± 0.0066 | 0.9506 ± 0.0055 |
| Clindamycin | Support Vector Machine | 0.8256 ± 0.0120 | 0.8150 ± 0.0128 |
| Tetracycline | LightGBM | 0.9115 ± 0.0080 | 0.9120 ± 0.0092 |
| Tigecycline | Support Vector Machine | 0.8136 ± 0.0413 | 0.2455 ± 0.1701 |
| Cotrimoxazole | LightGBM | 0.8863 ± 0.0075 | 0.7902 ± 0.0114 |
| Ciprofloxacin | LightGBM | 0.9543 ± 0.0032 | 0.9647 ± 0.0042 |
| Teicoplanin | Support Vector Machine | 0.8704 ± 0.0191 | 0.6529 ± 0.0809 |
| Rifampicin | LightGBM | 0.9193 ± 0.0107 | 0.7223 ± 0.0331 |
